# Gene family information facilitates variant interpretation and identification of disease-associated genes

**DOI:** 10.1101/159780

**Authors:** Dennis Lal, Patrick May, Kaitlin E. Samocha, Jack A. Kosmicki, Elise B. Robinson, Rikke S. Møller, Roland Krause, Peter Nüernberg, Sarah Weckhuysen, Peter De Jonghe, Renzo Guerrini, Lisa M. Neupert, Juliana Du, Eduardo Perez-Palma, Carla Marini, EuroEpinomics-RES Consortium, James S. Ware, Mitja Kurki, Padhraig Gormley, Sha Tang, Sitao Wu, Saskia Biskup, Annapura Poduri, Bernd A. Neubauer, Bobby P. Koeleman, Katherine L. Helbig, Yvonne G. Weber, Ingo Helbig, Amit R. Majithia, Aarno Palotie, Mark J. Daly

## Abstract

Differentiating risk-conferring from benign missense variants, and therefore optimal calculation of gene-variant burden, represent a major challenge in particular for rare and genetic heterogeneous disorders. While orthologous gene conservation is commonly employed in variant annotation, approximately 80% of known disease-associated genes are paralogs and belong to gene families. It has not been thoroughly investigated how gene family information can be utilized for disease gene discovery and variant interpretation. We developed a paralog conservation score to empirically evaluate whether paralog conserved or nonconserved sites of in-human paralogs are important for protein function. Using this score, we demonstrate that disease-associated missense variants are significantly enriched at paralog conserved sites across all disease groups and disease inheritance models tested. Next, we assessed whether gene family information could assist in discovering novel disease-associated genes. We subsequently developed a gene family *de novo* enrichment framework that identified 43 exome-wide enriched gene families including 98 *de novo* variant carrying genes in more than 10k neurodevelopmental disorder patients. 33 gene family enriched genes represent novel candidate genes which are brain expressed and variant constrained in neurodevelopmental disorders.

---

Identifying disease-causing mutations from the vast sea of benign rare variants in a genome continues to be an important challenge in both clinical and research settings despite the availability of large-scale population sequence references^1,2^. While functional studies would be the gold standard for determining causality, they are expensive, time consuming, and require extensive expert knowledge. Even in the case of coding variants, appropriate assays testing the disease-relevant impact of a variant on gene function are not readily available for the majority of genes and variants. Because of these limitations, *in silico* prediction tools have become the method of choice. These tools predict the deleteriousness of variants by assessing sequence conservation between species, structural constraints, amino acid physiochemical properties, and/or known annotations^3–5^. Sequence conservation, in which mammalian or vertebrate genomes are aligned to identify conserved segments among orthologs has been a component of nearly every such method, under the sensible premise that since genes often retain function through evolution, those genic elements that remain constant throughout evolution are more likely essential to gene function. The high average sequence similarity of homologs of disease-associated genes often translates into high conservation and therefore higher pathogenicity scores for *in silico* prediction tools based on vertical conservation profiles (Fig. 1 left).

Frequently overlooked, however, is the sequence conservation within species available from paralogous genes. Paralogs are defined as genes related via ancient duplication events^6–8^. Paralogous sequences are often utilized in alignments together with orthologous sequences in several variant prediction methods. However, the contribution of human paralogous sequences in variant annotation has not been thoroughly explored. Moreover, most of these methods rely on trained multi-feature classifier, which do not allow biological interpretation of results. Paralogs within gene families show often a relatively high sequence divergence. This greater variance among paralogs in comparison to orthologs, reflecting deeper evolutionary roots of some duplication events, suggests an unexplored opportunity that might aid in classifying residues as more or less functionally important (Fig. 1 right) based on their horizontal conservation profiles within gene families. The human genome harbors 3,348 protein-coding gene families, ranging from 2 to 46 paralog members and accounting for 72% of all human protein-coding genes (ENSEMBL v20150512). It has been shown that approximately 80% of disease-associated genes have paralogs in human^9^.

**Figure 1.**
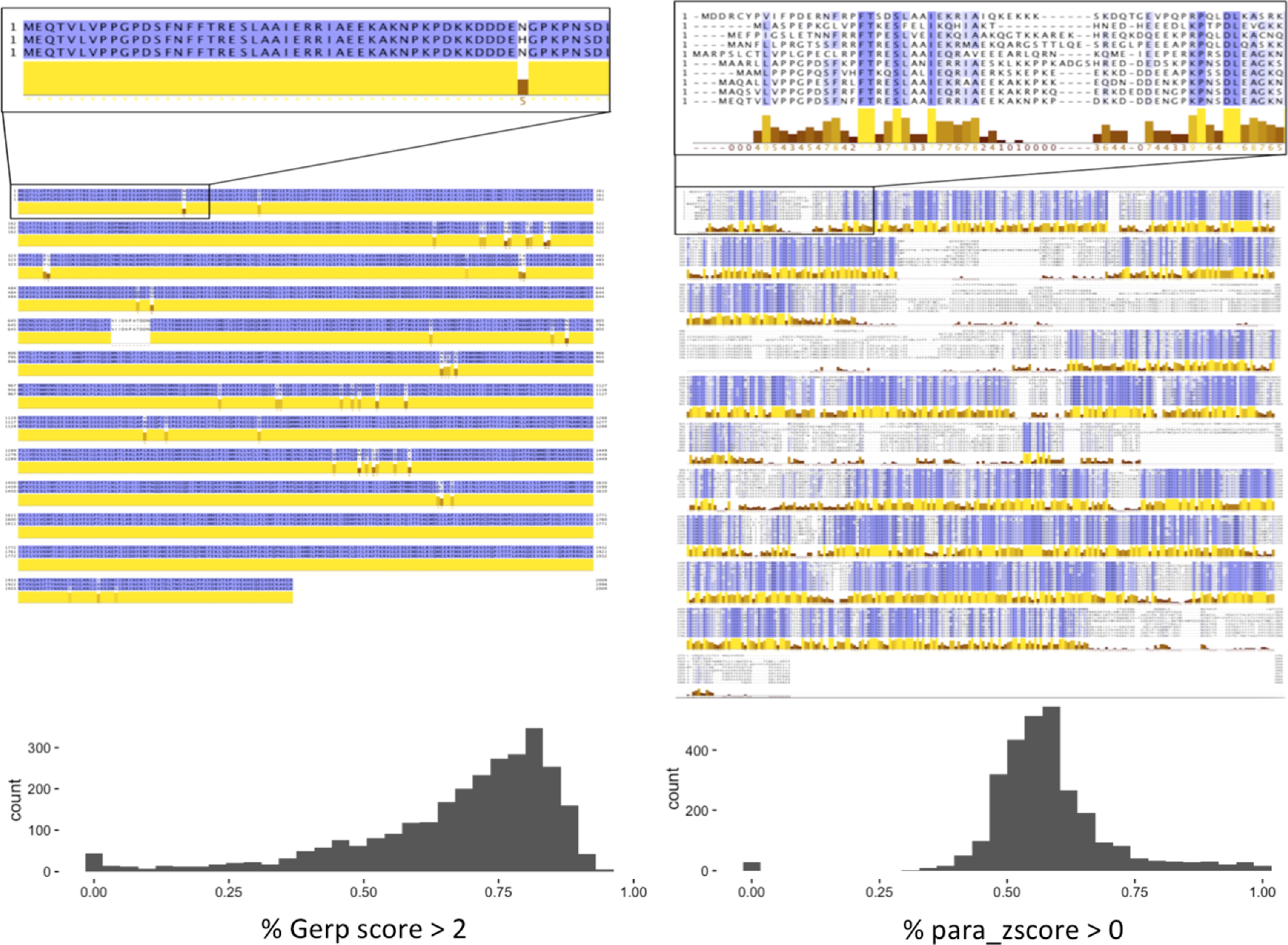
Vertical (ortholog) vs. horizontal (paralog) conservation. Top: Protein sequence alignment of voltage-gated sodium channels. **Top left**: Alignment of *Homo sapiens* (NP_001159435.1), *Bos taurus* (NP_001180147.1) and *Mus musculus* (NP_001300926.1) SCN1A protein sequences. High sequence similarity is depicted by violet amino acid coloring and yellow conservation bars below the alignment using Jalview. **Top right**: Protein alignment in Jalview of all members of the human voltage-gated sodium channel gene family (*SCN1A, SCN2A, SCN3A, SCN4A, SCN5A, SCN7A, SCN8A, SCN9A, SCN10A, SCN11A).* This alignment of paralogues shows less conservation compared to the *SCN1A* vertical cross species alignment on the left. **Bottom left**: GERP score analysis over all genes within gene families (homolog conservation is measured by % of all nucleotides per gene with GERP scores >2). **Bottom right**: Distribution % nucleotides per gene within gene families having para_zscores >0. Homolog conservation is generally much more uniform and homogeneous than paralog conservation.

The association of diseases with gene families has been well established in the last two decades. For example, protein kinases comprise one of the largest families of evolutionarily related proteins and >500 distinct kinases are encoded by ∼2% of all human genes. Variants in kinase genes have been found to underlie many human diseases, particularly developmental and metabolic disorders, as well as certain cancers^10^. Another example represents the family of keratin genes. Variants in these genes alter the structure of keratins, which prevent them from forming an effective network of keratin intermediate filaments. Without this network, cells become fragile and are easily damaged, making tissues less resistant to friction and minor trauma^11^. More recent large scale sequencing studies on neurodevelopmental disorders (NDDs) have independently identified multiple paralogous genes associated with the same or related NDD (e.g., the family of voltage-gated sodium channel genes: *SCN1A, SCN2A, SCN8A*; the family of chromodomain-helicase-DNA-binding proteins: *CHD2, CHD4,* CHD8^12–15^. This raises the question whether other genes within the same gene family are also associated with NDDs. Since in these studies truncating variants in paralogs often show consistent associations to NDDs, we sought to explore whether paralog information could refine our interpretation of missense variation. Paralogs often have similar, but not identical, protein sequences (Fig. 1 right) and amino acids conserved across all paralogs might well be pointing to essential residues critical for protein function. As such, variants changing paralog conserved amino acids may plausibly be more deleterious than variants changing amino acids in paralog non-conserved sites and therefore be more likely to confer risk to disease. Two previous studies have highlighted the utility of systematic functional annotation of disease-causing residues across human paralogs for genes associated with Long-QT-syndrome, Brugada syndrome, and catecholaminergic polymorphic ventricular tachycardia^16,17^. Both studies showed improved variant interpretation by comparing corresponding mutations in paralogs in different patients with the same phenotype. To our knowledge, it has not been empirically investigated on genome-wide scale whether disease associated missense variants reside in paralog conserved or non-conserved sites.

Statistical power for the discovery of disease associated genes is the greatest in genetically homogeneous patient groups. NDDs are phenotypically and genetically heterogeneous. Several of the NDD disease-associated genes are pleiotropic and appear in clinically distinct NDDs (sub-NDDs) indicating a shared molecular pathology. Even in NDD cohorts with >1000 trios, the majority of disease-associated genes have fewer than 10 *de novo* variants^12–15^. To increase the statistical power in genetic studies, gene set enrichment analysis is often applied to discover pathways associated with diseases of the same or a similar phenotype^18,19^. To date, the utility of gene families as gene sets for large-scale mutational burden analyses and candidate gene identification has not been systematically conducted.

In this manuscript, we evaluate and develop the use of ‘paralog conservation’ and provide evidence that this offers powerful addition to variant annotation over and above conventional conservation metrics and other annotations as follows: (1) We develop a novel paralog-based conservation score (“para_zscore”, see Online Methods and **Supplementary Fig. 1**) and demonstrate that paralogs highlight a smaller, more refined set of conserved residues than conventional ortholog conservation. (2) We further empirically demonstrate that disease-associated missense variants across all genetic disorders are strongly enriched at paralog-conserved sites. (3) We demonstrate that restricting analysis to missense variants at only paralog conserved sites increases the power to discover novel disease associated genes. (4) Leveraging these observations, we create a novel gene-family burden test of association and identify, using a cohort of 10,060 NDD trios, gene families with *de novo* variant burden including novel candidate disease genes.

## Results

### Allele frequency of paralog conserved and non-conserved sites in the general population

We first sought to test whether paralog conserved amino acids were more likely to confer strong disease risk when altered in comparison to sites not conserved in paralogs. As an initial test, we correlated the allele frequency of 1.3 million missense variants affecting genes with paralogous family members in 60,706 reference individuals from the Exome Aggregation Consortium^2^ (ExAC) with their paralog conservation measured by para_zscore. Testing 1,213,427 amino acids across all human gene families, we found that allele frequency decreases with increasing paralog conservation, indicating stronger evolutionary selection against variation in paralog conserved sites overall (Fig. 2 top left panel).

To further compare the utility of paralog versus ortholog amino acid conservation, we analyzed *de novo* missense variants identified in 10,068 NDD patients and 2,078 controls. Using the GERP score we measured the (vertical) divergence across 35 mammalian species^20^. Given that only a small fraction of patient variants is expected to be associated with disease, we did not expect a major shift between the patient and control variant distributions. We did however observe a more significant difference between variant scores in patients compared to controls using the para_zscore (p<10^−5^) than we saw for GERP (p=.006; Fig. 2 top right panel).

### Paralog conservation of missense *de novo* mutations in NDD patients and controls

To investigate the degree to which *de novo* mutations (DNMs) in NDDs are enriched in paralog-conserved segments, we compared the variant distribution of DNMs in 10,068 NDD patients to 2,078 individuals without NDDs. To increase the signal, we excluded those DNMs present in ExAC^2,21^. We compared the distributions of para_zscores for synonymous and missense variants (Fig. 2 bottom left) expecting only an enrichment for missense variants due to the fraction of variants associated with disease. We observe a significant shift towards paralog-conserved (*P*= 2.5 × 10^−6^) sites for missense DNM variants in NDD patients but not in controls (*P*=0.87).

**Figure 2.**
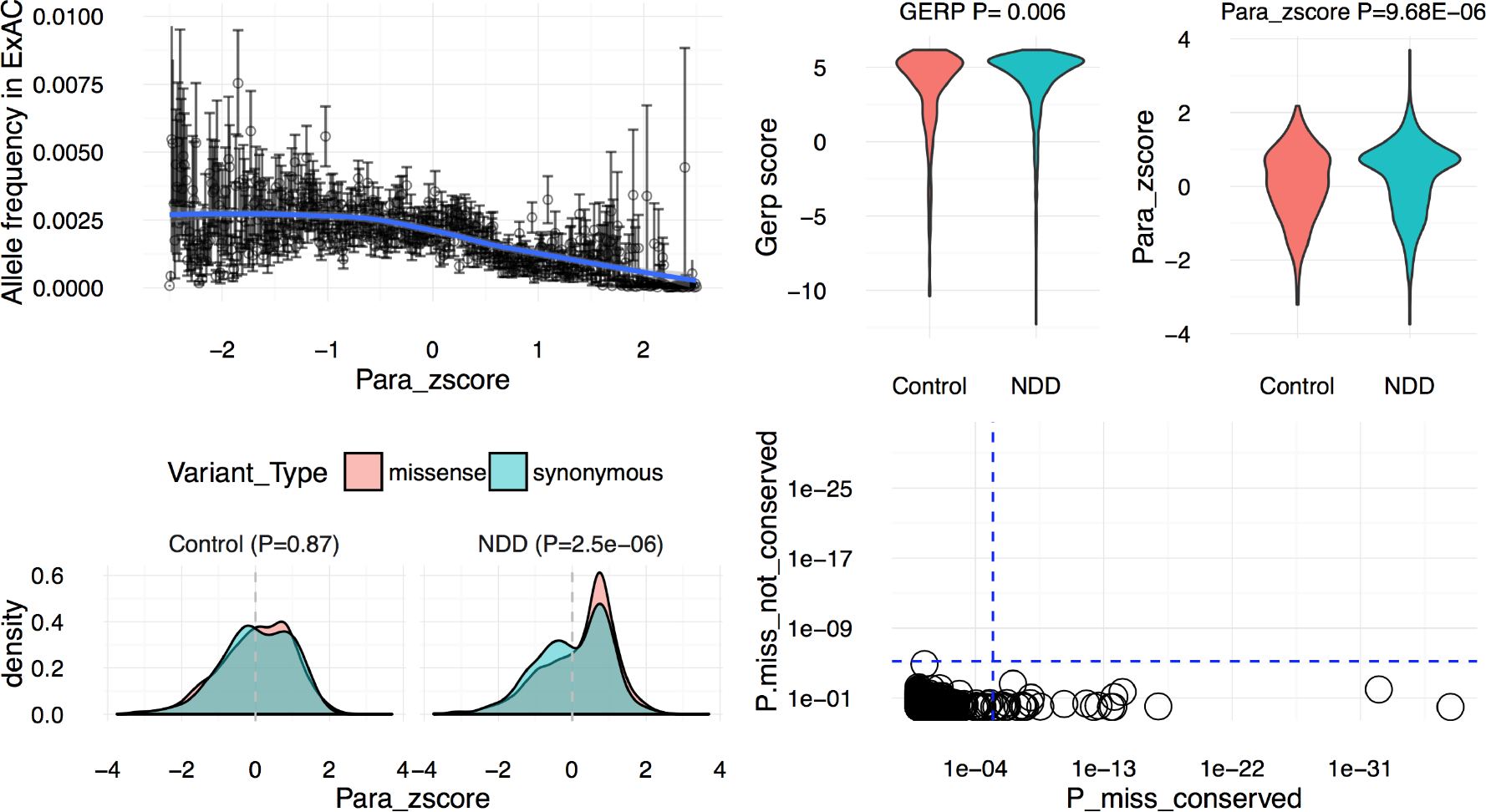
Assessment of paralog conservation. Top left. ExAC allele frequency vs. gene family (paralog) conservation (given as Z-score). The allele frequency of 1,213,427 missense variants decreases with increasing para_zscores for the tested variants. **Top right:** Ortholog vs. paralog conservation of missense variants. Ortholog conservation is measured by GERP scores, paralog conservation by para_zscores. Para_zscores better separate patient from control variants compared to GERP scores. Higher para_zscores are only slightly enriched in NDD patient variants compared to controls, which is in line with the expectation that not all patient *de novo* missense variants contribute to disease. **Bottom left:** Distribution of NDD patient and control missense and synonymous para_zscores depicted by density plots. Patient, but not control, missense variants are enriched at paralog conserved sites relative to the synonymous variant distribution. *P*-values were calculated using a Wilcoxon test. **Bottom right:** Identification of missense variant gene family enrichment in NDD patients for paralog conserved missense variants but not for the paralog non-conserved variants. NDD associated missense variants are enriched in paralog conserved sites. Y-axis: Missense variant enrichment analysis considering only paralog non-conserved sites across genes of each gene family (para_zcore ≤0). X-axis: Missense variant enrichment analysis considering only paralog conserved sites (para_zcore >0). None of the gene families show exome-wide significant enrichment for paralog non-conserved sites. 26 gene families (depicted by circles) show exome-wide significant *de novo* missense variant burden at paralog conserved sites. The significance threshold was calculated by Bonferroni correction for testing 5 × 2871 gene families (*P* = 3.48 × 10^−6^) and is depicted by the blue dotted line.

We evaluated the contribution of paralog conserved and non-conserved missense variants to NDDs. Considering 2,871 gene families encapsulating 9,991 genes, we observed 27 significantly enriched (p < 3.48 × 10^−6^) gene families in the patient cohort identified in an analysis of paralog conserved missense variants only, but none in the parallel, comparably powered, analysis of only non-conserved sites (Fig. 2 bottom right, **Supplementary Fig. 1B**). Although many of these genes also show burden for protein truncating variants (PTVs), the paralog enrichment is specific for missense variants since we did not identify a shift towards paralog conserved sites for nonsense variants (**Supplementary Fig. 2**). This is in line with expectations because nonsense-mediated-decay is the expected disease mechanism. Furthermore, missense variant enrichment at paralog conserved sites is not detectable in genes without DNM burden in this study (**Supplementary Fig. 2**).

### Enrichment of missense variants at paralog conserved sites across all diseases and associated genes

To investigate whether the enrichment for disease variants at paralog conserved sites is specific to our NDD cohort or can be generally observed across other disease categories, we extracted all missense variants from the ClinVar database (accessed June 2016) and classified the disorders into ICD chapters and sub-groups. We compared the paralog conservation for all gene families harboring disease variants against the missense variants in the same genes from ExAC. Overall, we detect a strong enrichment for disease variants at paralog-conserved sites compared to ExAC missense variants (*P* = 2.2 × 10^−287^, shift in para_zscore = 0.92, CI=0.87-0.98; Fig. 3). Our analyses indicate that disease-associated variants more often affect amino acids that are conserved across all members of the gene family. The shift towards paralog conserved sites for patient variants in comparison holds true for almost all disease groups tested.

**Figure 3.**
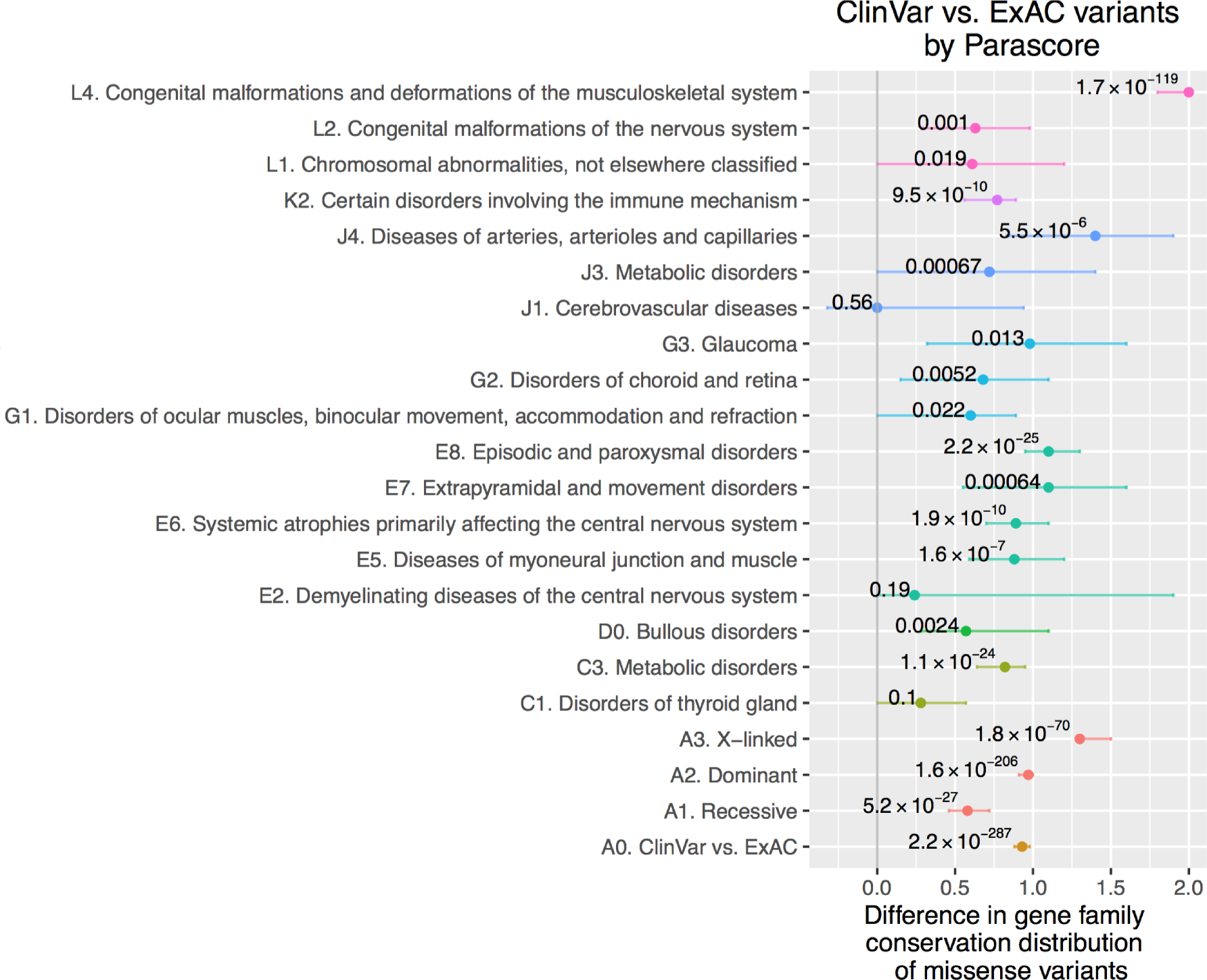
ClinVar disease vs. ExAC missense variant para_zscore distribution. We scored ClinVar variants and variants in a reference population (ExAC) for the same genes in a range of disorders and grouped these into ICD10 code defined disorder classes (see Online Methods). The distribution of ClinVar variants is shifted towards paralog conserved sites compared to missense variants in ExAC as depicted by estimated differences in location plotted in the x-axis for the vast majority of disorders tested. The shift to paralog conserved sites is depicted by estimates greater zero. *P*-values were calculated using a Wilcoxon test.

### Gene family enrichment in NDDs and sub-phenotypes

After establishing that disease-associated missense variants are enriched at paralog conserved sites (Fig. 2 bottom right, Fig. 3), we assessed the degree to which gene family information could assist in discovering novel disease-associated genes. We analyzed again our heterogeneous cohort of 10,068 NDD patients with autism spectrum disorder (ASD; N=3982), developmental delay (DD; N=5226) and epilepsy (EPI; N=822). We extended the approach of Samocha et al., 2014^22^, to gene families to identified gene families with significant enrichment of mutations in NDD patients. We included protein truncating variants (PTVs) across the whole sequence as well as missense variants at paralog conserved protein sites absent from ExAC in the analysis.

We identified 43 gene families (1.49% of all gene families) enriched for *de novo* paralog-conserved missense and PTVs (Bonferroni correction significance threshold for testing 5 × 2871 gene families = 3.48 × 10^−6^; Table 1). In all 43 gene families, the most frequently mutated gene and often additional genes harboring *de novo* variants are brain-expressed (**Supplementary Fig. 3**). Within the enriched gene families, 94 gene family members (paralogs) carried at least one DNM vs. 59 gene family members without DNMs. In total, 7.47% of all NDD patients carried a *de novo* paralog conserved missense or PTV in the 43 enriched gene families. In the NDD patients, we found 753 DNMs in 43 gene families while only 49.92 DNMs were expected (*P* << 1.0 × 10^−100^). The paralog conserved missense variant enrichment signal of these genes was 7.8-fold (observed DNMs: 261, expected DNMs: 33.01). There was no signal if we examine paralog non-conserved missense variants in this group of genes (observed DNMs: 41, expected DNMs: 31.03, **Supplementary Table 2**). No enrichment was observed in the 2,078 individuals without a NDD (observed DNMs: 5, expected DNMs: 10.34, *P* = 1.0). The majority of the frequently mutated genes have previously been established as disease-associated genes by demonstrating an exome-wide significant DNM burden in disease-specific single gene enrichment studies^1212–15,23^ (Table 1, highlighted in black and bold). When removing all the established disease genes (Table 1, bold and black, N=42) from the analysis, we still observe a 4.72-fold enrichment (observed: 162 DNMs in the 43 enriched genes families, expected 34.27, *P* = 6.10 × 10^−56^). This enrichment increases to 5.28-fold when we removed all non-brain-expressed genes from the 43 enriched genes families (28.71 DNM expected vs. 149 DM observed, *P* = 3.72 × 10^−57^).

**Table 1.** **43 significantly enriched gene families in the combined *de novo* paralog conserved missense and PTV analysis for 10,068 NDD trios.** Only enriched gene families significant after applying the Bonferroni significance threshold for testing 5 × 2,871 gene families (3.48 × 10^−6^) are included. Bold-highlighted genes are affected by DNM and the number of DNM is indicated inside the soft-brackets. Genes in red have not previously been reported as significantly enriched in exome-wide ASD, DD or EPI studies. The full list gene families with variants can be found in **Supplementary Table 1**.

| Gene families | DNMs expected | DNMs observed | P | 5264 DD patients | 3982 ASD patients | 822 EPI patients | 2087 controls |
| --- | --- | --- | --- | --- | --- | --- | --- |
| ARID1B (40), ARID1A (5) | 0.96 | 45 | 4.28E-58 | 39 | 6 | 0 | 0 |
| SCN2A (38), SCN1A (18), SCN8A (11), SCN3A (4), SCN11A (2), SCN9A (1), SCN5A, SCN7A, SCN4A, SCN10A | 6.16 | 74 | 1.77E-52 | 36 | 16 | 22 | 0 |
| DDX3X (35), DDX3Y | 0.49 | 35 | 8.94E-52 | 34 | 1 | 0 | 0 |
| DYRK1A (26), DYRK1B (1) | 0.37 | 27 | 1.89E-40 | 21 | 5 | 1 | 0 |
| EP300 (20), CREBBP (16) | 1.39 | 36 | 8.81E-38 | 30 | 5 | 1 | 1 |
| KCNQ2 (23), KCNQ3 (7), KCNQ5 (3) | 1.14 | 33 | 3.09E-36 | 27 | 2 | 4 | 0 |
| SYNGAP1 (24), DAB2IP, RASAL2 | 1.33 | 24 | 4.35E-22 | 17 | 6 | 1 | 0 |
| STXBP1 (20), STXBP3 (1), STXBP2 | 0.88 | 21 | 5.93E-22 | 14 | 2 | 5 | 0 |
| GRIN2B (16), GRIN2A (6), GRIN2C (1), GRIN2D | 1.89 | 23 | 1.46E-17 | 17 | 3 | 3 | 0 |
| CTNNB1 (16), JUP (1) | 0.79 | 17 | 2.27E-17 | 15 | 2 | 0 | 0 |
| CHD2 (16), CHD1 (3) | 1.15 | 19 | 4.00E-17 | 10 | 8 | 1 | 0 |
| PURA (13), PURB | 0.41 | 13 | 1.13E-15 | 12 | 0 | 1 | 0 |
| CHD3 (10), CHD4 (6), CHD5 (5) | 1.94 | 21 | 3.45E-15 | 19 | 2 | 0 | 0 |
| TCF4 (11), TCF12 (3), TCF3 (2) | 0.93 | 16 | 5.76E-15 | 14 | 2 | 0 | 1 |
| CDK13 (14), CDK12 | 0.66 | 14 | 1.71E-14 | 13 | 1 | 0 | 0 |
| PPP2R5D (16), PPP2R5A (1), PPP2R5B, PPP2R5C, PPP2R5E | 1.25 | 17 | 3.62E-14 | 16 | 1 | 0 | 1 |
| WDR45 (11), WDR45B | 0.32 | 11 | 7.60E-14 | 9 | 0 | 2 | 0 |
| MEF2C (11), MEF2D (2), MEF2A | 0.58 | 13 | 7.77E-14 | 9 | 2 | 2 | 0 |
| EHMT1 (13), EHMT2 | 0.66 | 13 | 3.93E-13 | 13 | 0 | 0 | 0 |
| **FOXP1 (10), **FOXP2 (4)**, FOXP4** | 0.86 | 14 | 6.02E-13 | 12 | 2 | 0 | 0 |
| **FOXG1 (11), FOXQ1, FOXN3, FOXN2** | 0.39 | 11 | 6.36E-13 | 7 | 1 | 3 | 0 |
| **CHD8 (12), **CHD7 (7), CHD9 (1)**** | 2.30 | 20 | 7.73E-13 | 11 | 9 | 0 | 0 |
| **CSNK2A1 (12), CSNK2A3, CSNK2A2** | 0.63 | 12 | 5.00E-12 | 10 | 1 | 1 | 0 |
| **CACNA1E (10), **CACNA1A (9), CACNA1B (1)**** | 2.72 | 20 | 1.53E-11 | 13 | 1 | 6 | 0 |
| **HDAC8 (9), **HDAC3 (2), HDAC1 (1), HDAC2**** | 0.83 | 12 | 1.06E-10 | 11 | 0 | 1 | 0 |
| **GNAO1 (7), GNAI1 (7), **GNAZ (1), GNA11 (1)**,**<br>**GNA14, GNAI3, GNAT2, GNAT3, GNAT1, GNAI2,**<br>**GNA15, GNAQ** | 1.82 | 16 | 1.31E-10 | 13 | 2 | 1 | 0 |
| **CASK (8), DLG4 (6), **DLG2 (1)**, DLG1, DLG3** | 1.58 | 15 | 1.67E-10 | 12 | 1 | 2 | 0 |
| **GATAD2B (10), GATAD2A** | 0.65 | 10 | 2.10E-09 | 10 | 0 | 0 | 0 |
| **MED13L (12), **MED13 (2)**** | 1.68 | 14 | 3.47E-09 | 10 | 4 | 0 | 0 |
| **TBL1XR1 (9), TBL1Y, TBL1X** | 0.52 | 9 | 5.01E-09 | 7 | 2 | 0 | 0 |
| **PHIP (10), BRWD3 (2)** | 1.46 | 13 | 5.64E-09 | 10 | 2 | 1 | 0 |
| **CTCF (9), CTCFL** | 0.56 | 9 | 9.00E-09 | 7 | 2 | 0 | 0 |
| **RAB11A (3), RAB2A (2), RAB11B (2), RAB14 (2),**<br>****RAB19 (1), RAB43 (1)**, RAB39A, RAB25, RAB4B,**<br>**RAB39B, RAB4A, RAB30, RAB2B** | 1.01 | 11 | 1.14E-08 | 9 | 2 | 0 | 0 |
| **GABRB3 (8), **GABRB2 (3), GLRA2 (1), GABRB1 (2), GLRB (1)**,**<br>**GLRA1, GABRR1, GABRD, GABRR3, GABRP, GLRA4, GABRQ, GABRR2,**<br>**GLRA3** | 2.35 | 15 | 3.15E-08 | 7 | 4 | 4 | 0 |
| **TCF20 (9), **RAI1 (3)**** | 1.40 | 12 | 3.33E-08 | 11 | 1 | 0 | 1 |
| **NFIX (6), **NFIA (2), NFIB (2)**, NFIC,** | 0.90 | 10 | 4.31E-08 | 8 | 2 | 0 | 1 |
| **USP9X (10), **USP24 (1)**, USP9Y** | 1.18 | 11 | 5.29E-08 | 9 | 2 | 0 | 0 |
| **SATB2 (9), SATB1** | 0.75 | 9 | 1.04E-07 | 9 | 0 | 0 | 0 |
| **HECW2 (9), HECW1 (1)** | 1.04 | 10 | 1.61E-07 | 8 | 1 | 1 | 0 |
| **SOX11 (4), SOX4 (3), **SOX9 (1), SOX10 (1)**, SOX17,**<br>**SOX1, SOX8, SOX7** | 0.82 | 9 | 2.21E-07 | 9 | 0 | 0 | 0 |
| **ZBTB18 (6), ZBTB3** | 0.28 | 6 | 5.43E-07 | 6 | 0 | 0 | 0 |
| **TCF7L2 (6), **TCF7L1 (1)**** | 0.53 | 7 | 1.48E-06 | 4 | 3 | 0 | 0 |
| **TBR1 (6), EOMES** | 0.35 | 6 | 2.01E-06 | 1 | 5 | 0 | 0 |

Several of these genes, previously not associated with any disease, represent likely NDD-associated genes. First, in one out of the 43 enriched gene families, we observe that the novel gene harbors more DNM variants than the established disease-associated gene of the same gene family (*CHD3* vs. *CHD4*; Table 1). Second, four families show genome-wide gene family enrichment without prior evidence for any gene on the genome-wide level, even though some individual genes are known disease genes, such as *CACNA1A* (**Supplementary Table 3**). These gene families include the (CACNA1[E/A/B] – family, RAB[2(A/B)/4(A/B)/11(A/B)/14/19/25/30/39(A/B)/43] - family, the HECW[1/2] - family, the SOX[1/4/7/8/9/10/11], TCF7L[1/2] - family and the TBR1/EOMES - family). To find further evidence of disease association for less frequently mutated gene family members, we systematically investigated the evolutionary variant intolerance^22^ (“constraint”) and brain expression levels for all mutated paralog genes within the enriched gene families (**Supplementary Fig. 3**). We observed that 33 paralog genes of the enriched gene families with DNMs are under evolutionary constraint and brain expressed (**Supplementary Table 3**), showing the same signature as the known disease genes in the same families (**Supplementary Fig. 3**). Although none of these genes have previously been reported to be significant on an exome-wide level, 57.57% (19/33) of the novel disease-associated genes have been previously reported in the literature as carrying a rare single nucleotide or copy number variant affecting the gene (**Supplementary Table 3**). For 63.63% (21/33) of the genes, available mice models show neurological and/or behavioral phenotypes supporting the disease association (**Supplementary Table 3**). Given these multiple lines of evidence, in addition to the sequence and expression pattern similarity to the known disease genes in the same families, we consider this list of 33 genes as highly promising candidate disease genes (**Supplementary Table 3**).

### Sub-disorder specificity of DNM enriched gene families

While the initial analysis considered all NDDs as a single group, we next explored whether the NDD sub-phenotypes (ASD, DD and EPI) contributed equally to the enrichment of the 43 gene families (Table 1) or if a specific sub-NDD contributes more to the enrichment and subsequently carries larger mutational burden than the other NDDs. Compared to single gene enrichment studies, DNM enriched gene families harbor larger numbers of variants due to the sum of mutations in several mutated paralogous genes, which increases testing power for investigating disease specificity. The increased number of variants per family permits frequency comparisons of likely functional related proteins across NDDs. We tested the 43 PTV and paralog-conserved missense enriched gene families (Table 1) to evaluate gene family specific sub-NDD burden. To avoid introducing a bias, we included all mutated genes of each gene family in the analysis, regardless of the PTV or missense intolerance and gene tissue expression analysis in **Supplementary Fig. 3**.

Overall, 31.91% of all *de novo* variants in patients with DD affected one gene in the 43 gene families, compared to 13.53% in the ASD cohort and 26.03% in the EPI cohort. Only two out of the 43 enriched gene families (Table 1) showed significant sub-disorder enrichment (either ASD, DD or EPI) after multiple testing correction. The DDX3[X/Y]-family is 20.55-fold enriched in DD patients in comparison with mutation frequencies in ASD and EPI patients combined (*P*=3.17 × 10^−6^). The voltage-gated sodium channels (SCN[1/2/3/4/5/7/8/9/10/11]A), have significantly higher mutational burden in the EPI cohort compared to variant frequencies in the ASD and DD cohorts combined (OR=4.96, *P*=6.06 × 10^−8^,) while representing the most frequent gene family with *de novo* mutations in all sub-NDDs (EPI=2.67%, DD=0.68%, ASD=0.40% carrier rate). Post-hoc phenotype analysis of the ASD cohort revealed that ASD patients with voltage-gated sodium channel mutations have a 5.08-fold higher frequency of seizures and 8.5-fold higher rate of low IQ compared to ASD patients without mutations in this gene family.

### Variant interpretation and correlation of para_zscore with an experimentally derived mutation intolerance score

The vast majority of current variant prediction methods are nucleotide based and do not allow visualization of scores across the protein sequence. Paralog conservation can identify stretches of conserved amino acids which can overlap functional domains; however not all annotated domains are paralog conserved and harbor disease variants (Fig. 4). The para_zscore is able to identify paralog conserved and non-conserved amino acid residues within known functional domains (Fig. 4) and thus is an illustrative tool to visualize the predicted mutation tolerance of a protein sequence across the amino acid sequence. As an example, we wanted to visualize para_zscore amino acid conservation alongside functionally derived data. Variants in *PPARG* cause Mendelian lipodystrophy and increase risk of type 2 diabetes (T2D). Recently, a complete prospective functional assessment of all possible 9,595 amino acid exchanges in *PPARG* was conducted^24^. The authors developed a pooled functional assay in human macrophages, experimentally evaluated all protein variants, and used the experimental data to train a variant classifier by supervised machine learning (“mut_tol”, Fig. 5). Comparing the experimental derived data with the *PPARG* para_zscores, we observed high correlation (*r*^2^=0.71, *P* = 4.08 × 10^−77^, Pearson’s product-moment correlation), supporting the usefulness of the para_zscore as a biological interpretable pathogenicity metric. In comparison, averaged mutation tolerance scores (see Online Methods) for Polyphen^5^, CADD^4^, SIFT^3^ and GERP^20^ show less correlation (*r*^2^= 0.28-0.56) with mut_tol (Fig. 5).

**Figure 4.**
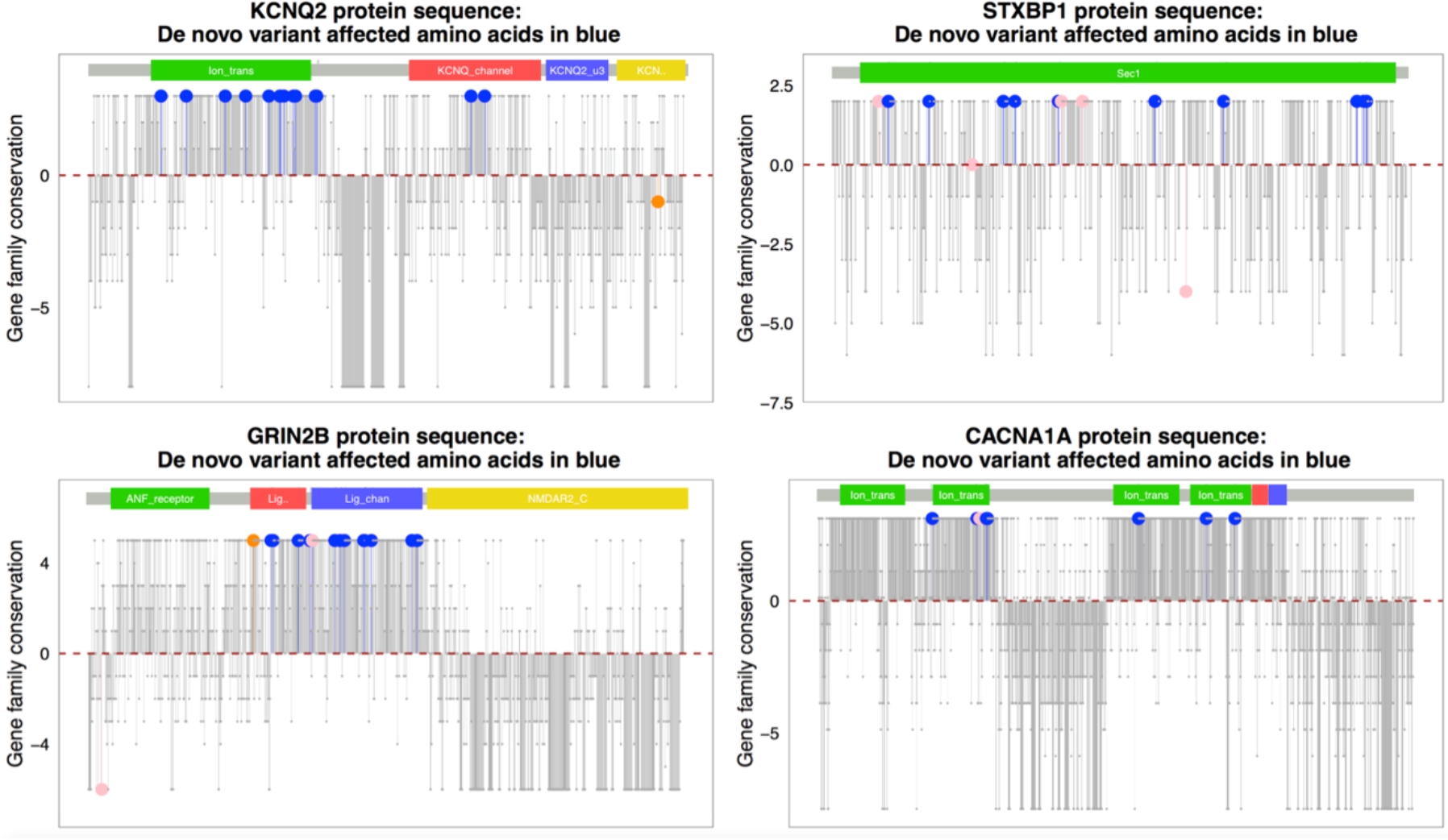
Visualization of para_zscores of proteins encoded by *KCNQ2, STXBP1,CACNA1A,* and *GRIN2B*. Protein sequence is plotted from left to right. Each bar and dot represents one amino acid. Amino acids affected by a missense mutation in the NDD cohort are colored blue, patient PTVs are depicted in pink, and synonymous variants in orange. Amino acid residues with no mutations are colored grey. Y-axis: Positive values indicate paralog conservation and the highest score indicates that these amino acids are identical over all gene family members. The red dotted lines indicate the mean paralog conservation of each protein sequence and bars below the mean indicate regions of low paralog conservation, thus higher sequence variability over all members of the gene family.

**Figure 5.**
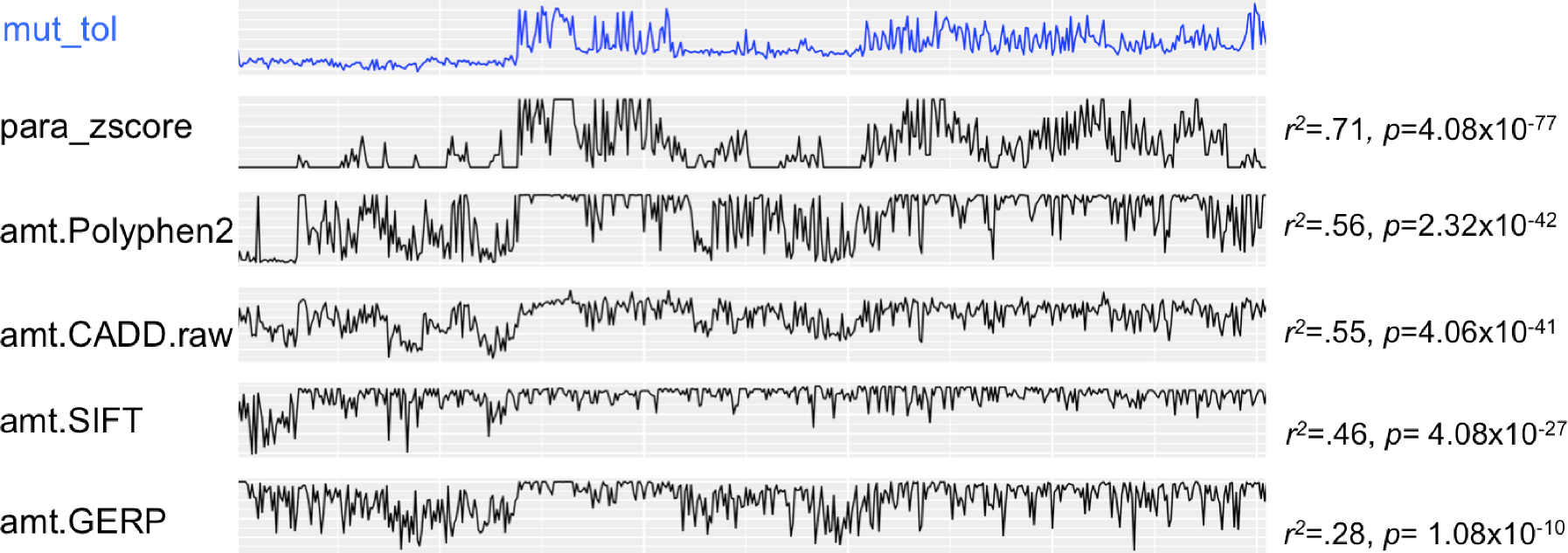
Comparison of experimentally derived mutation tolerance scores and para_zscores for *PPARG*. Raw *PPARG* function scores for each of the 9,595 amino acid exchanges plotted according to the amino acid position along the *PPARG* sequence. Functional scores summed by amino acid position are plotted to the right, showing the level of tolerance for any amino acid substitution differing from the human reference genome (derived from Majithia et al., 2016^24^). PPARG functionally generated mut_tol scores (blue), para_zscores and averaged mutation tolerance scores (amt) for PolyPhen2, CADD, and SIFT. All scores were correlated with the experimentally derived mut_tol score and *r* and *p*-values are shown next to the score distribution.

## Discussion

Sequence conservation across gene families has been extensively discussed in the literature. On the one hand, orthologous domain pairs tend to be significantly more structurally similar than paralogous pairs at the same level of sequence identity^25^. On the other hand, it has been shown that paralogs are functionally not necessarily redundant and the average fitness cost of loss of a paralogous gene is at least equal deleting single non-paralogous genes in yeast^26^. In addition, protein complexes have been documented to use paralog switching as a mechanism for the regulation of complex stoichiometry^27^. For example, many receptors in the human brain consist of multiple protein subunits, many of which have multiple paralogs and are differentially expressed across brain regions and developmental stages. The brain can tune the electrophysiological properties of synapses to regulate plasticity and information processing by switching from one protein variant to another. Such condition-dependent variant switch during development has been demonstrated in several neurotransmitter systems including NMDA and GABA^28^. Naturally, the question arises whether paralog conserved or non-conserved sites of the protein sequence are essential for function. Across multiple analyses in this study, we empirically demonstrate that disease-associated missense variants are enriched at paralog conserved sites and that this information can offer substantial value in mutation annotation on top of widely-used annotation methods.

We developed a gene family DNM enrichment framework and computed a novel amino acid paralog conservation score, the “para_zscore”, applicable to 42% of all genes in the human genome. Application of the genome-wide paralog conservation score demonstrates that NDD and pathogenic variants in ClinVar affect paralog conserved, but not non-conserved, sites when compared to mutations in controls and missense variants in ExAC^2^, the vast majority of which are presumed benign. It has recently been proposed that conserved residues within gene family members (paralogs) are under evolutionary constraint and that *in silico* annotations of known disease associated residues across families of related proteins can guide variant interpretation^17^. Our results support the idea that paralogs share a similar “core” molecular function of the ancestral gene, since variants in these sites are enriched for patient missense variants indicating hereby a reduction in evolutionary fitness. Evolutionary younger paralogs show higher functional redundancy^29^. To control the functional diversity within gene families and to increase within family sequence similarity, we built sub-groups for each gene family using pairwise alignment length cut-offs of >80% aligned amino acids^30^.

NDDs represent a genetic and phenotypic heterogeneous group of diseases for which pathogenic variants in individual disease genes are rare. Using a gene family version of a recently established DNM enrichment framework^22^ for 10,068 NDD patients, we identified 43 PTV and paralog conserved missense DNM enriched gene families. Besides highlighting four gene families with genome-wide gene family enrichment without carrying any previously exome-wide established disease gene (CACNA1[E/A/B] - family, RAB[2(A/B)/4(A/B)/11(A/B)/14/19/25/30/39(A/B)/43] - family, HECW[1/2] - family, SOX[1/4/7/8/9/10/11], TCF7L[1/2] - family, TBR1/EOMES - family), we additionally report 33 with first time statistical support as disease genes. Pathogenic variants in all of these genes are too rare to reach individual exome-wide significant enrichment. However, the individual genes belong to a gene family with an exome-wide significant enrichment, are brain expressed, and are under evolutionary constraint in the general population^2^. Notably, three of the 43 enriched genes belong to chromatin helicase DNA-binding proteins gene families. Besides the established genes, we observe also five novel candidate disease genes in this group (*CHD1, CHD3, CHD5, CHD7, CHD9)* with DNM in 20 patients in this study. All five genes represent valid candidates for association of NDDs based on our detailed analysis (for details see **Supplementary Table 3**). Chromatin remodeling is one of the mechanisms by which gene expression is regulated developmentally^31^, perhaps explaining the susceptibility to NDDs when mutated.

Given the larger number of DNMs in gene families compared to single genes, we investigated if any of the 43 enriched gene families are specifically enriched in patients within the EPI, ASD, or DD sub-cohorts. The vast majority of gene families did not show significant sub-NDD specificity. This phenomenon can be explained in several ways. Most importantly, there are overlapping phenotypes in the different studies. Patients within the developmental delay cohort include patients with ASD^15^. For NDD phenotype definitions overlap significantly, allowing for including of patients with combined phenotypes into either of the DD, ASD, or EPI cohort. However, we showed that paralog conserved missense and PTVs affecting the voltage-gated sodium alpha channel family are clearly enriched in patient within the EPI cohort, but the same class of variants can also be found at lower frequency in DD, and rarely in ASD. Analysis of the ASD patients with variants in voltage-gated sodium alpha channels indicated that the majority of the ASD patients with such variants had seizures. However, we were not able to investigate the DD patients for the seizure phenotype due to the lack of available phenotypic data. A higher degree of specificity of pathogenic variants in voltage-gated sodium channels in patients with epilepsy is supported biologically by the function of the channels. Lack of the sodium channel on inhibitory neurons leads to decreased inhibition and net excitation. This molecular pathology resembles the long-standing idea that epilepsy is a disorder based on imbalances between synaptic excitation and inhibition^32^.

The para_zscore has single value for each position; it is therefore possible to plot the score along the entire protein sequence (see Fig. 4). We propose that plotting the para_zscore is a useful tool as it visualizes likely essential protein regions of high paralog conservation, and thus intuitively supports decision making for variant prioritization (e.g., functional testing or drug target development). The scores are available at https://git-r3lab.uni.lu/genomeanalysis/paralogs. Supporting this utility, we demonstrated that the para_zscore exhibits high correlation (r^2^=0.71) with experimentally derived mutation tolerance scores across the entire *PPARG* protein sequence. Other generated averaged mutation tolerance scores for commonly used classifiers showed less correlation. Notably, the other prediction tools are not made for protein conservation scoring. We suggest that our protein wide paralog conservation score could potentially also be used in targeted drug development to improve the efficacy of therapy. Paralogs share similar protein sequences or structural features, e.g. similar binding pockets, e.g. a given compound may show an increased affinity to bind its paralogs, possibly resulting in unexpected cross-reactivity and undesired side effects. Using the paralog conservation score in drug target design could therefore rule out or reduce such cross-reactivity effects.

Although our results demonstrate the utility of paralog conservation, the ideal composition of the gene family and choice of protein isoform may differ depending on the individual research question. In addition, while conservation across orthologs or paralogs can be indicative of the necessary function of a given domain, absence of conservation does not *a priori* exclude functionally important domains within a protein. This consideration may be particularly relevant for diseases with a later onset that are less under evolutionary selection.

Overall, we provide extensive empirical evidence using multiple data sets (the entire ClinVar resource and published *de novo* variants from more than 10k NDD cases) that disease associated missense variants are enriched at paralog conserved sites. We demonstrate that integration of paralog conservation can be leveraged as powerful method for variant interpretation and discovery of new disease-associated genes. We provide a genome-wide "para_zscore" annotation file and pre-computed para_zscores for all human paralogs as individual files. This resource should enable data and molecular scientists to score and visualize variants/genes/proteins of interest and to integrate paralog conservation with existing variant annotation tools.

## Online Methods

### Patient and genetic data

We analyzed 10,068 neurodevelopmental disorder (NDD) trios (probands and their unaffected parents) including 3,982 autism (ASD), 5,226 developmental delay (DD) and 822 severe epilepsy (EPI) patients. The ASD cohort was derived from published studies^13,33^. The DD cohort combined published *de novo* variants from previous DD and ID studies due to overlap in cohort ascertainment^14,15,23,34,35^. The EPI cohort included published trio data sets (356/822 trios; 43%)^12,36^ as well as 466 (56% of the 822 trios) unpublished exome-wide *de novo* data. New exome sequencing trio data were collected in multiple centers, companies and consortia, including the University Clinic of Kiel, Germany, the University Clinic of Tübingen, Germany, the Boston Children’s Hospital, USA, the University of Antwerp, the RES consortia, the ESES consortia and the DESIRE Consortia, Ambry Genetics and CeGat.

As control data in several analyses in the study, we used 2,078 trios sequenced with the same technology as the ASD patient cohort. These controls are unaffected siblings of the ASD patients^13,33^.

To ensure uniformity in variant representation and annotation across published datasets and with respect to the ExAC reference database^2^, we created a standardized variant representation using a Python implementation of vt^37^ and reannotated all variants from the different datasets with ANNOVAR^38^ using the RefSeq and Ensembl gene annotations (2016Feb01), ClinVar^39^ (v20160302) and the dbNSFP v3.0^40^ databases for various functional prediction tools and conservation scores such as GERP^20^, CADD^4^, Polyphen2^41^, and SIFT^3^.

### Construction of the paralog conservation score

We restricted the analysis to within human paralogs to identify gene family burden and amino acids in proteins which are essential for function in humans. It has previously been shown that the human genome harbors essential genes for normal development which are not present in great apes. Such an example represents *SRGAP2,* which is essential for neocortex expansion^42^. Ensembl defines gene families based on maximum likelihood phylogenetic gene trees^43^ based on the longest translation annotated in CCDS^44^ for each gene. Paralogs are then defined as genes of the same species related by a duplication event (as an inner tree node). First, we downloaded the human paralog definitions using the Ensembl BioMart system^45^ representing each gene with an Ensembl gene identifier. The paralogs could be grouped into 3,584 gene families. Ensembl IDs were then converted to HGNC gene names (http://www.genenames.org). Noncoding genes and genes without a HGNC symbol were excluded and only gene families with at least two HGNC genes were used for further analysis. CCDS data were downloaded the same day as HGNC and Ensembl data (v20150512).

In total, 3,348 gene families were defined with 13,570 HGNC genes. 1,815 families contained three or more paralogs. Next, we extracted the longest transcript from CCDS for each HGNC gene and constructed for each gene family a FASTA formatted file for multiple sequence alignment with MUSCLE^46^ including all paralog protein sequences. Evolutionary younger paralogs show higher functional redundancy^29^. To avoid alignments of strongly diverging sequences and to increase overall similarity, we built sub-groups for each gene family using pairwise alignment length cut-offs of >80% aligned amino acids^30^. The clusters (sub-groups) were defined by connected components within an alignment similarity graph in which two genes with >80% aligned residues were connected through an edge. Clusters were then defined as connected components within the graph. Only clusters with at least two proteins were further processed. In total, we generated 2,871 (sub) gene families comprising 8,233 genes. Each subgroup was re-aligned using MUSCLE. The MUSCLE output was then processed as input for Jalview^47^ to generate and extract conservation scores for each alignment position. The conservation score calculation in Jalview is based on the AMAS method of multiple sequence alignment analysis^48^. Conservation is measured here as a numerical index reflecting the conservation of physicochemical properties in the alignment. Identities score highest, and the next most conserved group contains substitutions to amino acids lying in the same physicochemical class. For each HGNC CCDS gene, the conservation scores at each position were extracted from the Jalview. Finally, to identify amino acids of high and low paralog conservation and make scores comparable between genes, the mean and the standard deviation conservation score over all amino acids per gene were calculated to compute a paralog conservation z-score (“para_zscore”) per amino acid position by subtracting the mean from the original score dividing the difference by the standard deviation. (**Supplementary Fig. 4**).

### Gene family enrichment analysis

To identify gene families with significant mutational burden, we adopted a *de novo* expectation model^22^ to assess mutation rates for nonsense, frameshift, or canonical splice disruptions (collectively termed protein truncating variants ‘PTVs’) and missense variants for gene families (missense+PTV). We derived gene-based rates of *de novo* mutation from local sequence context and summed the expectations and the observed counts for all members within each gene family^2^. The expected and observed numbers of *de novo* mutations in each variant class for NDD combined were compared using a Poisson distribution. Notably, the discovery of *de novo* burden in a gene family is more challenging compared to the single gene analysis because of larger amount of expected mutations due combining the expectations from all gene family members (including those which are not expressed in the tissue of interest, e.g., brain). We used a Bonferroni corrected significance threshold for the 2,871 gene families tested (<0.05). Furthermore, to exclude “passenger” variants and enrich for true disease variants we excluded *de novo* variants also seen in adult individuals without early onset NDDs in ExAC^2^ (n=60,706 exomes) prior to the enrichment analysis. Variants absent from this reference panel, which is a proxy for standing variation in the human population, are more likely to be deleterious. This conservative filter reduces the power for gene association discovery changes, but increases the plausibility for association for the identified variant with severe sporadic disorders.

### Sub-NDD enrichment analysis

Sub-NDD enrichment was calculated for the significantly enriched gene families in the paralog conserved missense and PTV combined analysis. We used Fisher’s exact test to compare the sub-NDD of interest (e.g., EPI) with the other sub-NDDs (e.g., ASD+DD).

### Paralog conserved site vs. paralog non-conserved sites missense enrichment analysis

Similar to the missense+PTV enrichment analysis, we adopted the *de novo* expectation model^22^ to assess mutation rates for missense variants only. In the missense enrichment analysis, we consider only missense variants and missense expectations for each gene family. We classified every amino acid position within each gene into ‘conserved sites’ with paralog conservation (para_zscore > 0, amino acids with higher paralog conservation than gene-specific mean) and ‘non-conserved sites’ without paralog conservation (para_zscore ≤ 0) and summed the observations for both groups independently across the family. The expected missense mutation rates were adjusted for the size of the paralog conserved sites and non-conserved sites, respectively. The observed variant counts were assigned to either of these two groups depending on the paralog conservation state of the mutated amino acid residue for all members within each gene family (**Supplementary Figure 1**). The expected and observed numbers of *de novo* mutations in each variant class for NDD combined was compared using a Poisson distribution. To exclude “passenger” variants and enrich for disease variants, we excluded *de novo* variants present as standing variation in the 60,706 individuals in ExAC^2^ prior to the enrichment analysis.

### Identification of brain expressed genes and evolutionary constraint

We extracted brain expression data from the Genotype-Tissue Expression consortia^49^ (GTEx) data and considered genes with >1 read per kilobase of transcript per million mapped reads (RPKM) in brain tissues as “ brain-expressed”. Gene loss-of-function intolerance (pLI) scores and gene missense intolerance scores were derived from ExAC^2^. We considered genes with missense Z scores > 3.09 or pLI scores ≥ 0.9 as intolerant of variants. Genes were classified as plausible novel disease genes for NDD if they were present in an exome-wide enriched gene family, brain expressed, and under constraint (either missense or PTV).

### Paralog conservation analysis of ClinVar variants vs. ExAC variants

The ClinVar database (v20160302)^39^ was filtered for missense variants that were classified as pathogenic. We excluded variants that were classified as pathogenic by one submitter and not pathogenic by another submitter. Next, we homogenized disease names by removing ambiguous subtype classifications. We removed variants present in ExAC^2^ to increase the quality of truly disease associated variants. Finally, we counted the number of variants in genes associated with a specific disease, and kept only disease-gene association supported by at least ten variants. 67 disorders remained in the analysis. For each disorder-gene combination, missense variants in the same gene were included as comparison in the data set. Next, we classified and grouped all 67 disorders by ICD10 codes, scored ClinVar and ExAC variants with the paralog score, and compared the paralog variant distribution using the Wilcoxon test (wilcox.test implemented in R). Notably, in nearly every ICD-10-CM category there is a sub-category named “other specified” condition (http://www.icd10monitor.com). Codes titled “other” or “other specified” are for use when the information in the medical record provides detail for which a specific code does not exist (https://www.cdc.gov). Due to the limited definition and corresponding limited interpretation of these heterogeneous groups of disorders we removed these from the analysis.

### *PPARG* mutation tolerance scores for para_zscore, GERP, CADD, PolyPhen2, and SIFT

The mutation tolerance score (see **Figure 5**) as defined in Majithia et al., 2016^24^, is given for every position of the protein sequence. Since the para_zscore is defined for every position of the protein, it can directly be used as a mutation tolerance score. For PolyPhen2 and SIFT^3^ we used the matrix of all possible amino acid exchanges represented as a *nx20* matrix (*n*=protein length; n=505 for *PPARG).* The exchange matrices for both scores were generated using the respective webtools (see URLs) using all amino acid changes as input. Then, for every possible amino acid substitution in hg19 the average score of all possible amino acid exchanges at a given position was calculated, e.g., if for a given codon *n* missense mutations could cause *m* different amino acid exchanges, we calculated the average score over the *m* possible amino acid exchanges as the mutation tolerance score at a given position. Since CADD^4^ and GERP^20^ scores are defined for every genomic position, we used for every possible amino acid exchange the average respective score over all possible nucleotide exchanges within a given codon resulting into the same amino acid exchange.

### Correlation between para_zscore and PPARG mut_tolerance score

The authors provided us with the experimentally derived and trained mutational tolerance score (“mut_tol”) for *PPARG.* A detailed description of the score can be found in Majithia et al., 2016^24^. In short, the authors used highly parallel oligonucleotide synthesis to construct a library encoding all 9,595 (505 positions × 19 amino acid changes at each position) possible single amino acid substitutions to prospectively characterize *PPARG* variants. They developed a pooled functional assay in human macrophages, experimentally evaluated all protein variants, and used the experimental data to train a variant classifier by supervised machine learning. We used the *cor {stats}* function in R^50^ for the correlation analysis of both scores.

## Data availability

The code for generating the underlying alignments and the para_zscore, additional code and examples are available under https://git-r3lab.uni.lu/genomeanalysis/paralogs. Additional data and annotations for the hg19 missense variants are available from: https://zenodo.org/record/817898. All *de novo* variants used in the analysis are listed in **Supplementary Table 4**.

## URLs

PolyPhen2, http://genetics.bwh.harvard.edu/pph2/; SIFT, http://sift.jcvi.org/www/SIFT_seq_submit2.html; GTEx portal, http://www.gtexportal.org/home/; ExAC, http://exac.broadinstitute.org; ANNOVAR, http://annovar.openbioinformatics.org; ClinVar, https://www.ncbi.nlm.nih.gov/clinvar; Ensembl biomart, www.ensembl.org/biomart; Ensembl compara, www.ensembl.org/info/genome/compara; HGNC, http://www.genenames.org; CCDS, https://www.ncbi.nlm.nih.gov/projects/CCDS.

## Acknowledgements

We thank Shamil Sunyaev for scientific exchange and comments on the evolutionary aspects of our results. Parts of the computational analysis was performed on the high-performance computer system of the University of Luxembourg. We thank Mungo Carstairs and Jim Procter from the Jalview team at the University of Dundee for their help with scripting Jalview. PM was supported by the JPND-Courage and FNR NCER-PD grants. RG was supported by the EU seventh Framework Programme (FP7) under the project DESIRE grant N602531 (to RG). DL received funds from the German Academic Exchange Service (DAAD), grant number 57073880.BAN received funding by the Deutsche Forschungsgemeinschaft (Ne 416/5-1).

## Author contributions

DL and PM performed the analyses. DL, PM, AP and MJD designed the experiment. DL and PM wrote the code. AP and MJD supervised the research. DL, PM, AP and MJD wrote the paper. KES, JAK, EBR, RSM, RK, PN, SW, PDJ, RG, LMN, JD, EPP, CM, JSW, MK, PG, ST, SW, SB, AP, BN, BPK, KLH, YGW, IH, ARM generated data. DL was responsible for the remainder. All authors revised and approved the final manuscript.

## Competing Financial Interests

ST, SW, and KLH are full time employees of Ambry Genetics.

## References

1. 1000 Genomes Project Consortium et al. A global reference for human genetic variation. Nature 526, 68–74 (2015).

2. Lek, M. et al. Analysis of protein-coding genetic variation in 60,706 humans. Nature 536, 285–291 (2016).

3. Kumar, P., Henikoff, S. & Ng, P. C. Predicting the effects of coding non-synonymous variants on protein function using the SIFT algorithm. Nat. Protoc. 4, 1073–1081 (2009).

4. Kircher, M. et al. A general framework for estimating the relative pathogenicity of human genetic variants. Nat. Genet. 46, 310–315 (2014).

5. Adzhubei, I. A. et al. A method and server for predicting damaging missense mutations. Nat. Methods 7, 248–249 (2010).

6. Soukup, S. W. Evolution by gene duplication. S. Ohno. Springer-Verlag, New York. 1970. 160pp. Teratology 9, 250–251 (1974).

7. Koonin, E. V. Orthologs, Paralogs, and Evolutionary Genomics. Annu. Rev. Genet. 39, 309–338 (2005).

8. Fitch, W. M. Distinguishing Homologous from Analogous Proteins. Syst.Zool. 19, 99 (1970).

9. Dickerson, J. E. & Robertson, D. L. On the Origins of Mendelian Disease Genes in Man: The Impact of Gene Duplication. Mol. Biol. Evol. 29, 61–69 (2012).

10. Lahiry, P., Torkamani, A., Schork, N. J. & Hegele, R. A. Kinase mutations in human disease: interpreting genotype-phenotype relationships. Nat. Rev. Genet. 11, 60–74 (2010).

11. Szeverenyi, I. et al. The Human Intermediate Filament Database: comprehensive information on a gene family involved in many human diseases. Hum. Mutat. 29, 351–360 (2008).

12. Allen, A. S. et al. De novo mutations in epileptic encephalopathies. Nature 501, 217–221 (2013).

13. De Rubeis, S. et al. Synaptic, transcriptional and chromatin genes disrupted in autism. Nature 515, 209–215 (2014).

14. Fitzgerald, T. W. et al. Large-scale discovery of novel genetic causes of developmental disorders. Nature 519, 223–228 (2014).

15. McRae, J. F. et al. Prevalence and architecture of de novo mutations in developmental disorders. Nature 542, 433–438 (2017).

16. Ware, J. S., Walsh, R., Cunningham, F., Birney, E. & Cook, S. A. Paralogous annotation of disease-causing variants in long QT syndrome genes. Hum. Mutat. 33, 1188–1191 (2012).

17. Walsh, R., Peters, N. S., Cook, S. A. & Ware, J. S. Paralogue annotation identifies novel pathogenic variants in patients with Brugada syndrome and catecholaminergic polymorphic ventricular tachycardia. J. Med. Genet. 51, 35–44 (2014).

18. Fromer, M. et al. De novo mutations in schizophrenia implicate synaptic networks. Nature 506, 179–184 (2014).

19. Homsy, J. et al. De novo mutations in congenital heart disease with neurodevelopmental and other congenital anomalies. Science 350, 1262–1266 (2015).

20. Goode, D. L. et al. Evolutionary constraint facilitates interpretation of genetic variation in resequenced human genomes. Genome Res. 20, 301–310 (2010).

21. Kosmicki, J. A. et al. Refining the role of de novo protein-truncating variants in neurodevelopmental disorders by using population reference samples. Nat. Genet. 49, 504–510 (2017).

22. Samocha, K. E. et al. A framework for the interpretation of de novo mutation in human disease. Nat. Genet. 46, 944–950 (2014).

23. Lelieveld, S. H. et al. Meta-analysis of 2,104 trios provides support for 10 new genes for intellectual disability. Nat. Neurosci. 19, 1194–1196 (2016).

24. Majithia, A. R. et al. Prospective functional classification of all possible missense variants in PPARG. Nat. Genet. 48, 1570–1575 (2016).

25. Peterson, M. E. et al. Evolutionary constraints on structural similarity in orthologs and paralogs. Protein Sci. 18, 1306–1315 (2009).

26. DeLuna, A. et al. Exposing the fitness contribution of duplicated genes. Nat. Genet. 40, 676–681 (2008).

27. Ori, A. et al. Spatiotemporal variation of mammalian protein complex stoichiometries. Genome Biol. 17, (2016).

28. Bar-Shira, O., Maor, R. & Chechik, G. Gene Expression Switching of Receptor Subunits in Human Brain Development. PLOS Comput. Biol. 11, e1004559 (2015).

29. Chen, S., Krinsky, B. H. & Long, M. New genes as drivers of phenotypic evolution. Nat. Rev. Genet. 14, 645–660 (2013).

30. Dufayard, J.-F. et al. Tree pattern matching in phylogenetic trees: automatic search for orthologs or paralogs in homologous gene sequence databases. Bioinforma. Oxf. Engl. 21, 2596–2603 (2005).

31. Thompson, P. M., Gotoh, T., Kok, M., White, P. S. & Brodeur, G. M. CHD5, a new member of the chromodomain gene family, is preferentially expressed in the nervous system. Oncogene 22, 1002–1011 (2003).

32. An Introduction to Epilepsy. (American Epilepsy Society, 2006).

33. Iossifov, I. et al. The contribution of de novo coding mutations to autism spectrum disorder. Nature 515, 216–221 (2014).

34. de Ligt, J. et al. Diagnostic Exome Sequencing in Persons with Severe Intellectual Disability. N. Engl. J. Med. 367, 1921–1929 (2012).

35. Rauch, A. et al. Range of genetic mutations associated with severe non-syndromic sporadic intellectual disability: an exome sequencing study. TheLancet 380, 1674–1682 (2012).

36. Appenzeller, S. et al. De Novo Mutations in Synaptic Transmission Genes Including DNM1 Cause Epileptic Encephalopathies. Am. J. Hum. Genet. 95, 360–370 (2014).

37. Tan, A., Abecasis, G. R. & Kang, H. M. Unified representation of genetic variants. Bioinformatics 31, 2202–2204 (2015).

38. Wang, K., Li, M. & Hakonarson, H. ANNOVAR: Functional annotation of genetic variants from high-throughput sequencing data. Nucleic Acids Res. 38, 1–7 (2010).

39. Landrum, M. J. et al. ClinVar: public archive of interpretations of clinically relevant variants. Nucleic Acids Res. 44, D862–868 (2016).

40. Liu, X., Wu, C., Li, C. & Boerwinkle, E. dbNSFP v3.0: A One-Stop Database of Functional Predictions and Annotations for Human Nonsynonymous and Splice-Site SNVs. Hum. Mutat. 37, 235–241 (2016).

41. Adzhubei, I., Jordan, D. M. & Sunyaev, S. R. Predicting Functional Effect of Human Missense Mutations Using PolyPhen-2. in Current Protocols in Human Genetics eds.Haines, J. L.et al. 7.20.1–7.20.41 (John Wiley & Sons, Inc., 2013). doi:10.1002/0471142905.hg0720s76

42. Dennis, M. Y. et al. Evolution of Human-Specific Neural SRGAP2 Genes by Incomplete Segmental Duplication. Cell 149, 912–922 (2012).

43. Vilella, A. J. et al. EnsemblCompara GeneTrees: Complete, duplication-aware phylogenetic trees in vertebrates. Genome Res. 19, 327–335 (2008).

44. Farrell, C. M. et al. Current status and new features of the Consensus Coding Sequence database. Nucleic Acids Res. 42, D865–D872 (2014).

45. Kinsella, R. J. et al. Ensembl BioMarts: a hub for data retrieval across taxonomic space. Database J. Biol. Databases Curation 2011, bar030 (2011).

46. Edgar, R. C. MUSCLE: multiple sequence alignment with high accuracy and high throughput. Nucleic Acids Res. 32, 1792–1797 (2004).

47. Waterhouse, A. M., Procter, J. B., Martin, D. M. A., Clamp, M. & Barton, G. J. Jalview Version 2––a multiple sequence alignment editor and analysis workbench. Bioinformatics 25, 1189–1191 (2009).

48. Livingstone, C. D. & Barton, G. J. Protein sequence alignments: a strategy for the hierarchical analysis of residue conservation. Bioinformatics 9, 745–756 (1993).

49. Lonsdale, J. et al. The Genotype-Tissue Expression (GTEx) project. Nat. Genet. 45, 580–585 (2013).

50. R Core Team. R: A language and environment for statistical computing. Vienna, Austria; 2014. URL http://www.R-Proj.org (2015).

